# Unveiling Novel Molecular Drivers in Breast Cancer Brain Metastasis: Multi-Omics Integration Identifies Downregulation of VCAN and Emerging Roles of ASCL2/GRAMD1A as Prognostic Biomarkers and Therapeutic Vulnerabilities

**DOI:** 10.1101/2025.08.18.670830

**Authors:** Hadith Rastad, Mahin Seifi-Alan, Azadeh Jalilian, Sanaz Seifi-Alan, Sedighe Hoseini, Haniyeh Rashidi, Kobra Hosseini, Mahnaz Seifi-Alan

**Author notes:** Correspondence to: Mahnaz Seifi-Alan, Non-Communicable Diseases Research Center, Alborz University of Medical Sciences, Karaj, Iran.

## Abstract

**Purpose:** Breast cancer brain metastasis (BCBM) presents a major clinical challenge, driven by molecular mechanisms that remain poorly characterized.

**Patients and methods:** Three RNA-seq datasets (GSE110590, GSE193103, GSE209998) were analyzed to identify BCBM-associated genes. Survival outcomes (2,976 primary tumors) were assessed via Kaplan-Meier (KM Plotter), genetic alterations via cBioPortal, pathways/networks via GeneMANIA/SIGNOR, and miRNA-mRNA interactions via miRNet. Drug candidates were prioritized using the CTD.

**Results:** TNFRSF9 and VCAN were downregulated (log2FC: −1.18 to −2.63), while GRAMD1A, ASCL2, TACC3, and PFKFB4 were upregulated (log2FC: +1.02 to +1.70). High PFKFB4 (HR=1.71) and TACC3 (HR=1.46) predicted poor survival, with VCAN suppression (Fold change (Fc) =0.24) and GRAMD1A elevation (Fc=1.31) confirmed in metastases. Pathways included ECM remodeling (VCAN), metabolic rewiring (PFKFB4), and mitotic instability (TACC3). miR-210-3p (hypoxia) and miR-27a-3p (angiogenesis) drove BCBM, countered by miR-335/34a. Drug candidates: Valproic Acid (TACC3/ASCL2), Vorinostat (VCAN), and CDK4/6 inhibitors.

**Conclusion:** This study identifies TNFRSF9, VCAN, GRAMD1A, ASCL2, TACC3, and PFKFB4 as key drivers of BCBM, with dysregulation linked to immune evasion, metabolic adaptation, and mitotic instability. Prioritized miRNAs (e.g., miR-210-3p) and repurposed drugs (e.g., Valproic Acid, Vorinostat) offer actionable therapeutic strategies. These findings advance precision approaches for BCBM, pending preclinical validation to translate targets into clinical practice.

## Introduction

Breast cancer brain metastasis (BCBM) is associated with poor prognosis, with only a small fraction living five years or longer. [1] Although systemic therapies have improved, the molecular adaptations enabling tumor cells to thrive in the brain’s nutrient-poor, immune-restricted milieu remain elusive. [2, 3] Current research prioritizes pan-metastatic drivers (e.g., invasion genes), overlooking brain-specific mechanisms. [4, 5] Disparate analyses of genomic, transcriptomic, and regulatory layers further obscure the identification of brain-specific therapeutic targets. Additionally, the absence of biomarkers capable of predicting BCBM risk or guiding targeted therapies highlights a pressing unmet need in precision oncology. [6–8]

To resolve these limitations, we conducted a multi-omics investigation integrating transcriptomic, genetic, and regulatory network analyses. Our strategy harmonized three independent BCBM cohorts to systematically differentiate brain-specific metastatic drivers from genes involved in general cancer progression. Upstream regulators, including miRNAs, were prioritized, and therapeutic vulnerabilities were assessed through computational drug repurposing. To our knowledge, this study is the first to unify genetic alterations, survival-associated expression patterns, pathway dysregulation, and miRNA-mRNA networks into a cohesive BCBM framework. By bridging mechanistic insights with clinical translatability, this work seeks to redefine BCBM biology and accelerate the development of targeted therapies for this lethal complication.

## Methods

### Differential Expression Analysis

To identify genes with altered expression in brain metastases, we analyzed three independent RNA-seq datasets (GSE110590, GSE193103, GSE209998) comparing primary breast tumors to metastatic lesions in the brain. Raw count matrices were processed through DESeq2 (v1.38.3) in R (v4.2.2) to pinpoint differentially expressed genes (DEGs). The full code for this analysis can be obtained from the corresponding author upon request.

For the GSE193103 and GSE209998 datasets, we implemented a two-stage workflow:

1. **Tissue-Specific Gene Exclusion**: By comparing normal brain and breast tissues (adjusted *p* < 0.05), we identified and removed genes with organ-specific expression to reduce confounding signals. In contrast, the GSE110590 dataset lacked matched normal tissue data, preventing tissue-specific filtering.
2. **Primary vs. Metastasis Comparison**: Brain metastasis samples were directly compared to primary breast tumors using a ∼Group design matrix, with primary tumors designated as the baseline

We applied likelihood ratio tests with independent filtering and stabilized log2 fold changes (LFC) using the ashr method to refine effect size estimates. DEGs were defined as genes with adjusted *p* < 0.05 and |LFC| >1 (a 2-fold change threshold).

### Intersection Analysis and Gene Prioritization

To isolate consistent BCBM-associated genes, we overlapped DEGs from all three datasets using a Venn diagram approach, identifying 56 consensus genes. From these, six genes were prioritized for deeper investigation. Selection criteria included:

- **Consistency**: Robust differential expression (significant *p* and LFC) across all datasets,
- **Novelty**: Underexplored roles in brain metastasis biology,
- **Translational Potential**: Biomarker suitability or druggability.

### Genetic Alteration Profiling

We interrogated genetic alterations in the six DEGs using cBioPortal (https://www.cbioportal.org/). Metastatic cohorts included Metastatic Breast Cancer (DFCI, Cancer Discov 2020), Metastatic Breast Cancer (INSERM, PLoS Med 2016), The Metastatic Breast Cancer Project (Archived, 2020), and The Metastatic Breast Cancer Project (Provisional, December 2021), while primary cohorts derived from following datasets Breast Cancer (METABRIC, Nature 2012 & Nat Commun 2016), Breast Invasive Carcinoma (British Columbia, Nature 2012), Breast Invasive Carcinoma (Broad, Nature 2012), Breast Invasive Carcinoma (Broad, Nature 2012), and Breast Invasive Carcinoma (TCGA, Firehose Legacy). Copy-number alterations (CNAs) were detected via GISTIC 2.0 and DNAcopy, with amplification/deletion frequencies calculated as the percentage of samples harboring alterations per gene. RNA-seq expression trends (e.g., **A**SCL2 upregulation in metastases) were cross-referenced with dominant CNAs (amplifications for upregulated genes, deletions for downregulated) to infer drivers of metastatic adaptation.

### Kaplan-Meier Survival Analysis

Using the KM Plotter tool (https://kmplot.com/analysis), we evaluated the prognostic relevance of the six DEGs in 2,976 primary breast tumors. Patients were stratified into “high” and “low” expression groups via the tool’s Auto Select Best Cutoff feature, which algorithmically identifies thresholds that maximize survival differences. The analysis included data from up to 10 years (120 months) of follow-up. Statistical significance was assessed with log-rank tests (p<0.05 threshold), and Cox proportional hazards models calculated hazard ratios with corresponding 95% confidence intervals. This linked BCBM-associated expression patterns (e.g., PFKFB4 upregulation) to survival outcomes in early-stage disease.

### Gene Expression Analysis via TNMplot

We assessed expression dynamics of the six DEGs across breast cancer progression using TNMplot (https://tnmplot.com/analysis/). RNA-seq data from 113 normal tissues, 1,097 primary tumors, and 7 metastatic lesions were analyzed. Median expression values and fold changes (Fc) were computed for tumor/normal and metastatic/tumor comparisons. Non-parametric Kruskal-Wallis tests identified global expression differences (*p* < 0.05), followed by Dunn’s post-hoc tests with Benjamini-Hochberg correction for pairwise group comparisons. Box plots visualized expression distributions, prioritizing non-parametric methods to account for non-normal data and limited metastatic samples.

### Pathway and Network Analysis

- **SIGNOR Database (**https://signor.uniroma2.it/): The six DEGs were mapped to curated signaling pathways in SIGnaling Network Open Resource (SIGNOR). Genes lacking entries (TNFRSF9, ASCL2, GRAMD1A) were flagged (Supplementary Figure 1), while interactions for PFKFB4, TACC3, and VCAN were summarized.
- **GeneMANIA (**http://genemania.org/): Interaction networks for downregulated (VCAN, TNFRSF9) and upregulated (PFKFB4, ASCL2, TACC3, GRAMD1A) genes were built using co-expression, physical interactions, and pathway data (max 20 interactions/gene).

### Functional Annotation and Hub Gene Identification

1. **Functional Annotation for DEGs**: Enrichr (https://maayanlab.cloud/Enrichr) performed pathway/ontology analyses for the six DEGs using Reactome, KEGG, and MSigDB Hallmark.
2. Hub Gene Workflow:
  ○ **Co-expression Networks**: TCGA breast cancer data (UALCAN (https://ualcan.path.uab.edu/analysis.html)) identified genes positively correlated with the six DEGs (Pearson *r* > 0.4). For ASCL2 and GRAMD1A, STRING v12 generated PPI networks directly due to sparse co-expression partners.
  ○ **PPI Networks**: STRING v12 (confidence score > 0.7) constructed interaction networks (Supplementary Figure 2).
  ○ **Hub Gene Ranking**: Cytoscape 3.10.3 with CytoHubba ranked genes by interaction degree, selecting the top 10 hubs per DEG (Supplementary Figure 3A).
  ○ **Enrichment Visualization**: DAVID [9, 10] performed GO/KEGG enrichment for hub genes (adjusted *p* < 0.05), visualized via bubble/bar plots using SRplot (https://www.bioinformatics.com.cn/) [11] (Supplementary Figure 3B&C).

### miRNA and Drug Prioritization

**miRNA Regulation**: miRNet 2.0 (https://www.mirnet.ca/miRNet/home.xhtml) predicted miRNAs targeting the six DEGs, filtered for breast cancer relevance. miRPathDB (https://mpd.bioinf.uni-sb.de/) linked prioritized miRNAs to metastasis pathways (hypoxia, ECM remodeling). miRCancer (http://mircancer.ecu.edu/search.jsp) validated roles (oncogenic/tumor-suppressive), and miRNA expression was confirmed in RNA-seq data (|log2FC| ≥0.5, adj. *p* < 0.05).

**Therapeutic Candidates**: The Comparative Toxicogenomics Database (CTD) (https://ctdbase.org/) identified drugs targeting DEGs, excluding environmental toxins and non-FDA compounds without clinical trial support. Priority was given to drugs with multiple CTD references or mechanistic relevance to metastasis (e.g., blood-brain barrier penetration for brain metastasis).

**Identification of Differentially Regulated Genes in Breast Cancer Brain Metastasis** We identified six genes—TNFRSF9, VCAN, GRAMD1A, ASCL2, TACC3, and PFKFB4— that showed consistent dysregulation across three transcriptomic datasets (GSE110590, GSE193103, GSE209998) when comparing brain metastases to primary breast tumors (Table 1). All genes met stringent statistical thresholds (unadjusted *p* < 0.01, adjusted *p* < 0.05) with uniform directionality. TNFRSF9 and VCAN were markedly suppressed in metastases (log2FC: −1.18 to −2.56), while GRAMD1A, ASCL2, TACC3, and PFKFB4 were robustly elevated (log2FC: +1.02 to +1.70). For instance, VCAN exhibited the strongest downregulation (log2FC: −2.63 in GSE110590, adjusted *p* = 5.46×10[[), whereas PFKFB4 showed the highest upregulation (log2FC: +1.70 in GSE193103, adjusted *p* = 1.74×10[[).

**Table 1:**
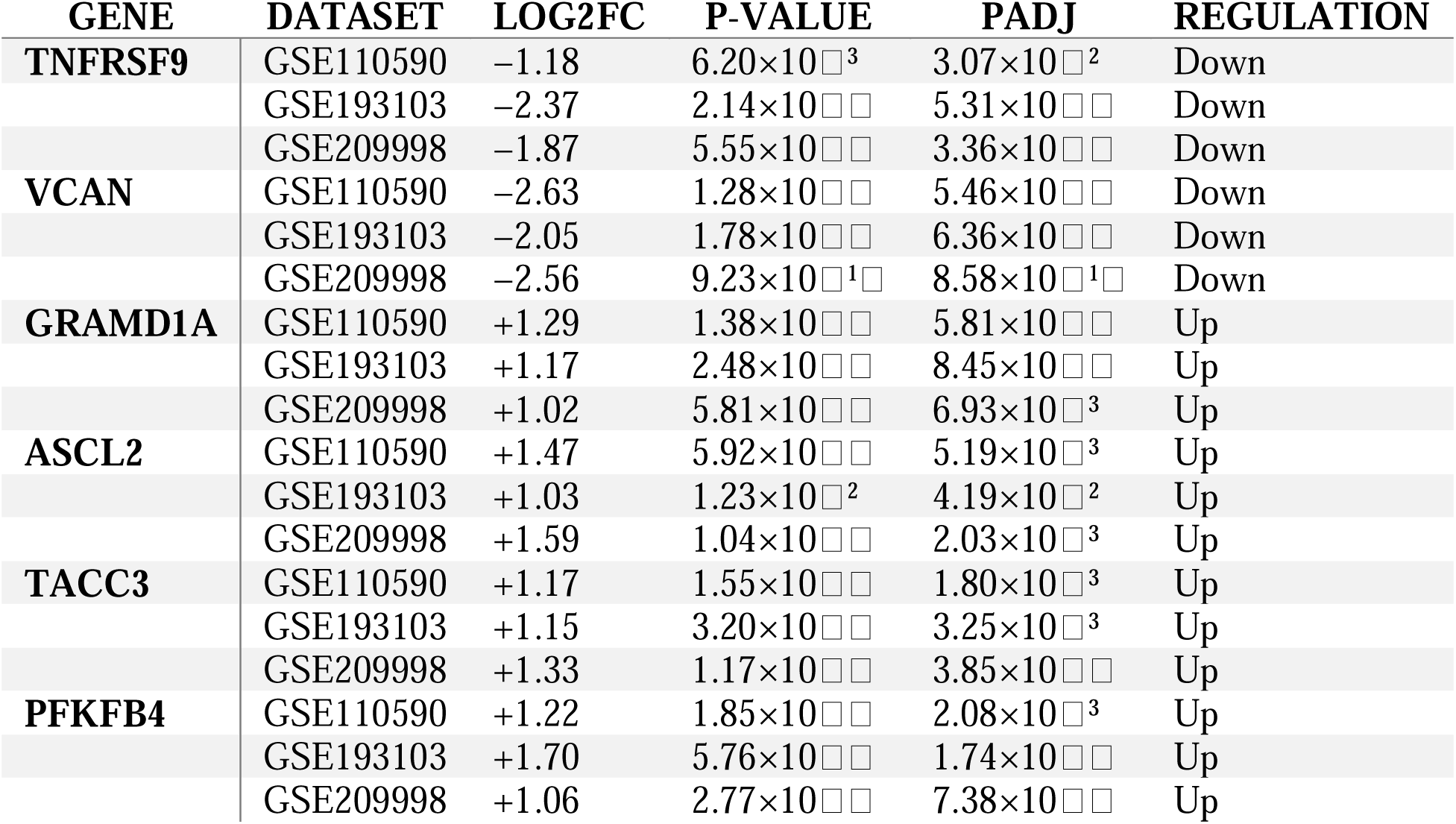
Consistent Dysregulation of Six Genes in BCBM Across Cohorts.

### Comparative Analysis of Genetic Alterations in Metastatic vs. Primary Breast Cancer

Metastatic tumors harbored distinct genetic alteration patterns compared to primaries, mirroring RNA-seq trends (Figure 1). Structural alterations were more frequent in metastases compared to primary tumors. Among upregulated genes (ASCL2, TACC3, PFKFB4, GRAMD1A), these alterations predominantly involved amplifications. Conversely, downregulated genes (VCAN, TNFRSF9) exhibited higher deletion rates in metastases relative to primary tumors. For example, VCAN’s 10-fold increase in metastatic alterations stemmed predominantly from deletions, directly linking genetic loss to its functional silencing. Similarly, TNFRSF9’s 12-fold alteration surge in metastases was driven by deletions, reinforcing its suppression. These findings highlight how metastatic progression hinges on *gain-of-function* amplifications (activating oncogenic pathways) and *loss-of-* function *deletions* (silencing metastasis suppressors).

**Figure 1.**
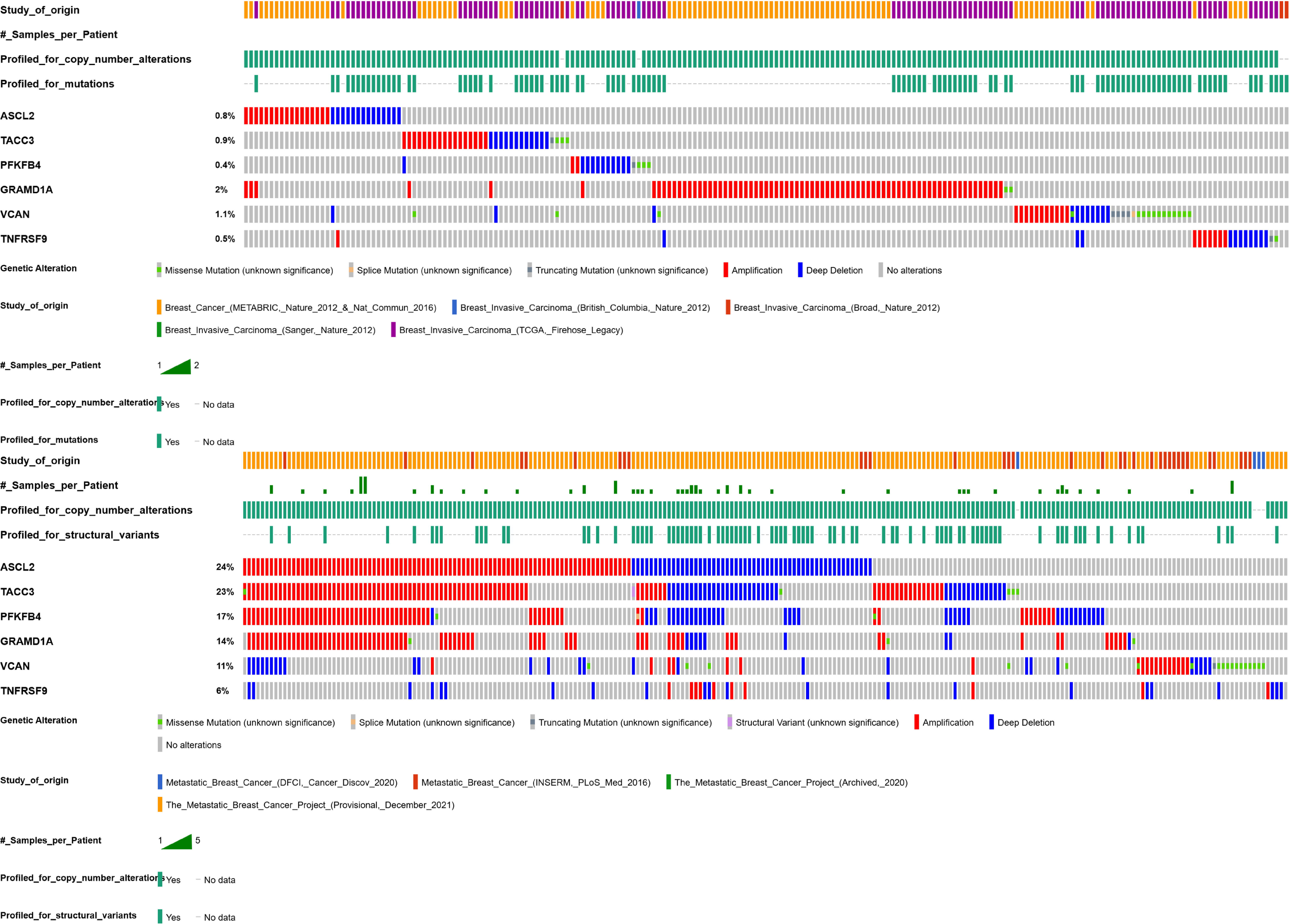
Analysis of genetic alterations related to the six DEGs across the genomic data sets (oncoprint) (**A**) from breast cancer invasive data, Altered in 203 (5%) of 3878 samples.; and **B**) from breast cancer metastasis dataset, Altered in 233 (40%) of 578 samples

### Survival Outcomes of Brain Metastasis-Associated DEGs in Primary Breast Tumors

Kaplan-Meier survival analysis revealed a striking inverse relationship between metastatic gene expression patterns and primary tumor outcomes (Figure 2). TNFRSF9 and VCAN— downregulated in metastases—correlated with improved survival when highly expressed in primaries (VCAN: HR = 0.77, *p* = 0.032). Conversely, upregulated genes PFKFB4, ASCL2, TACC3, and GRAMD1A predicted poor survival in primaries (PFKFB4: HR = 1.71, *p* = 1.7×10[[; TACC3: HR = 1.46, *p* = 0.0015). This alignment suggests these genes mark aggressive metastatic potential early in disease progression, functioning as biomarkers of outcome and progression.

**Figure 2.**
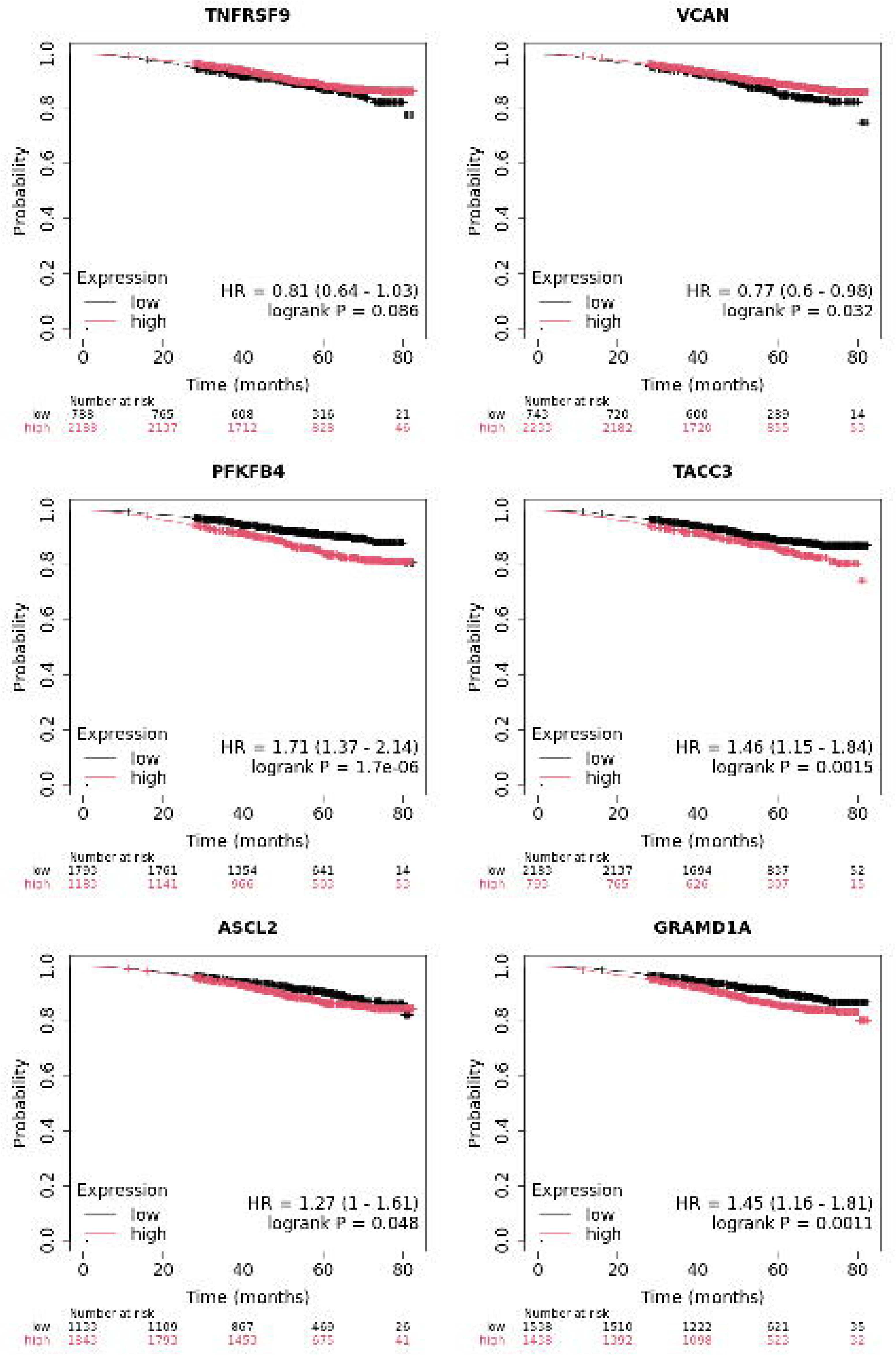
Kaplan-Meier overall survival (OS) curves for six differentially expressed genes (DEGs) in primary breast tumors. Genes were stratified into high-(red line) and low-expression (blue line) groups using the *KM Plotter* tool’s “Auto Select Best Cutoff” feature. Hazard ratios (HR) and log-rank *P*-values are displayed for each gene. **TNFRSF9** and **VCAN**, downregulated in brain metastases compared to primary tumors, were associated with improved survival when highly expressed in primary lesions. Conversely, **PFKFB4**, **ASCL2**, **TACC3**, and **GRAMD1A**, upregulated in brain metastases, correlated with significantly poorer survival when highly expressed in primary tumors. The “Number at risk” tables below each curve indicate cohort sizes at sequential follow-up intervals (months). This analysis links gene dysregulation patterns in brain metastases (loss of TNFRSF9/VCAN and gain of PFKFB4/ASCL2/TACC3/GRAMD1A) to divergent survival outcomes in primary breast cancer, highlighting their prognostic relevance. Data source: Breast Cancer RNA-seq cohort

### Differential Expression of Metastasis-Associated Genes in Breast Cancer: TNMplot Analysis

TNMplot analysis confirmed sustained dysregulation of these genes across cancer progression (Figure 2 & Table 2). VCAN showed marked suppression in metastases versus primaries (Fc = 0.24, *p* = 6.75×10[[), while TNFRSF9’s metastatic expression remained unchanged (Fc = 1.02, *p* = 0.238). Upregulated genes maintained elevated expression in metastases (e.g., GRAMD1A: median metastatic = 2,435 vs. tumor = 1,857), though metastatic/tumor differences lacked statistical significance (*p* > 0.05). (Table 2)

**Table 2:**
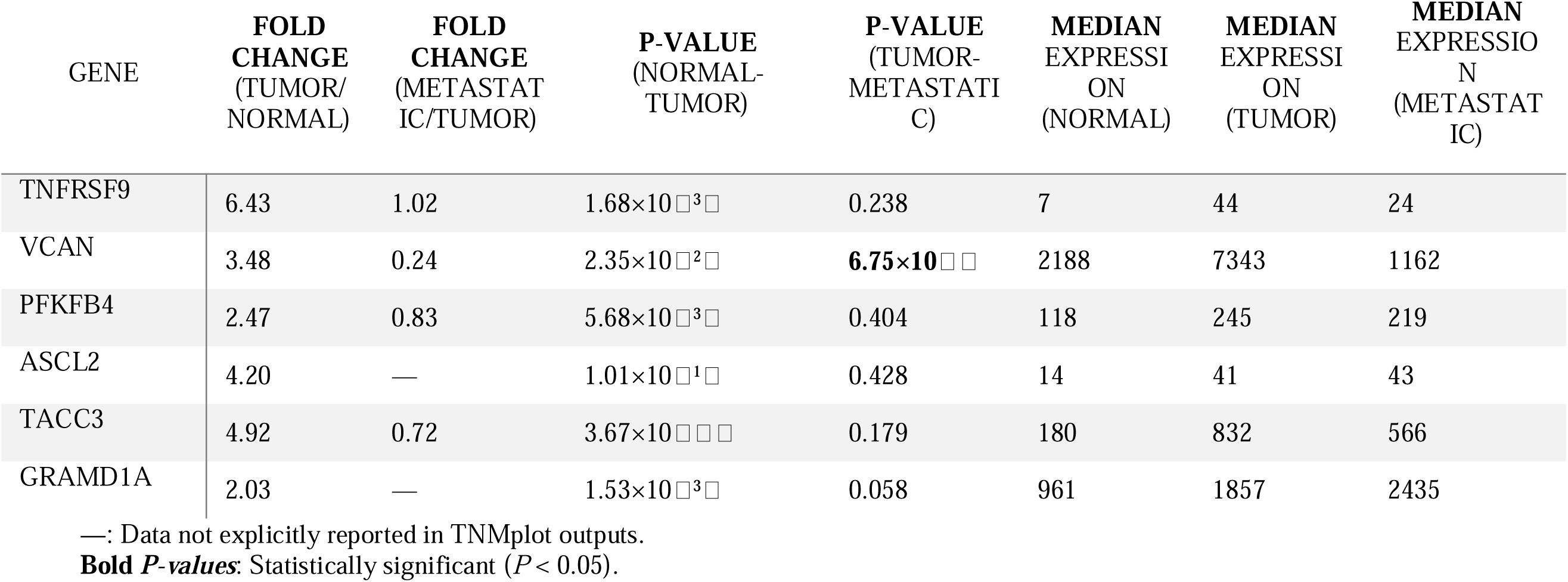
Expression Patterns of Six DEGs in Breast Cancer Progression based on TNMplot outputs.

### Sigaling Relations of Differentially Expressed Genes

SIGNOR database analysis mapped PFKFB4 to metabolic reprogramming (fructose-2,6-bisphosphate downregulation; PMID 30553771) and estrogen signaling via NCOA3 phosphorylation (PMID 29615789). VCAN was linked to p53-mediated ECM remodeling (PMID 12438652), while the AURKA-TACC3 axis showed high-confidence mitotic interactions (PMID 17545617). Absence of TNFRSF9, ASCL2, and GRAMD1A from SIGNOR underscores their non-canonical roles in metastasis

### GeneMANIA Network Analysis Reveals Functional Clusters

GeneMANIA networks segregated downregulated and upregulated genes into distinct functional modules (Figure 4 A&B). Downregulated genes (VCAN, TNFRSF9) clustered with immune regulators (*CXCL10*, *CCL8*) and ECM remodelers (*DSE*, *BCAN*), while upregulated genes (TACC3, PFKFB4) linked to metabolic (*PFKFB1*) and mitotic (*CDC25B*) pathways. VCAN’s interaction with *TLR6* and *CD44* suggests roles in immune evasion, whereas TACC3’s ties to *RPS6KB1* implicate mTOR-driven proliferation.

**Figure 3:**
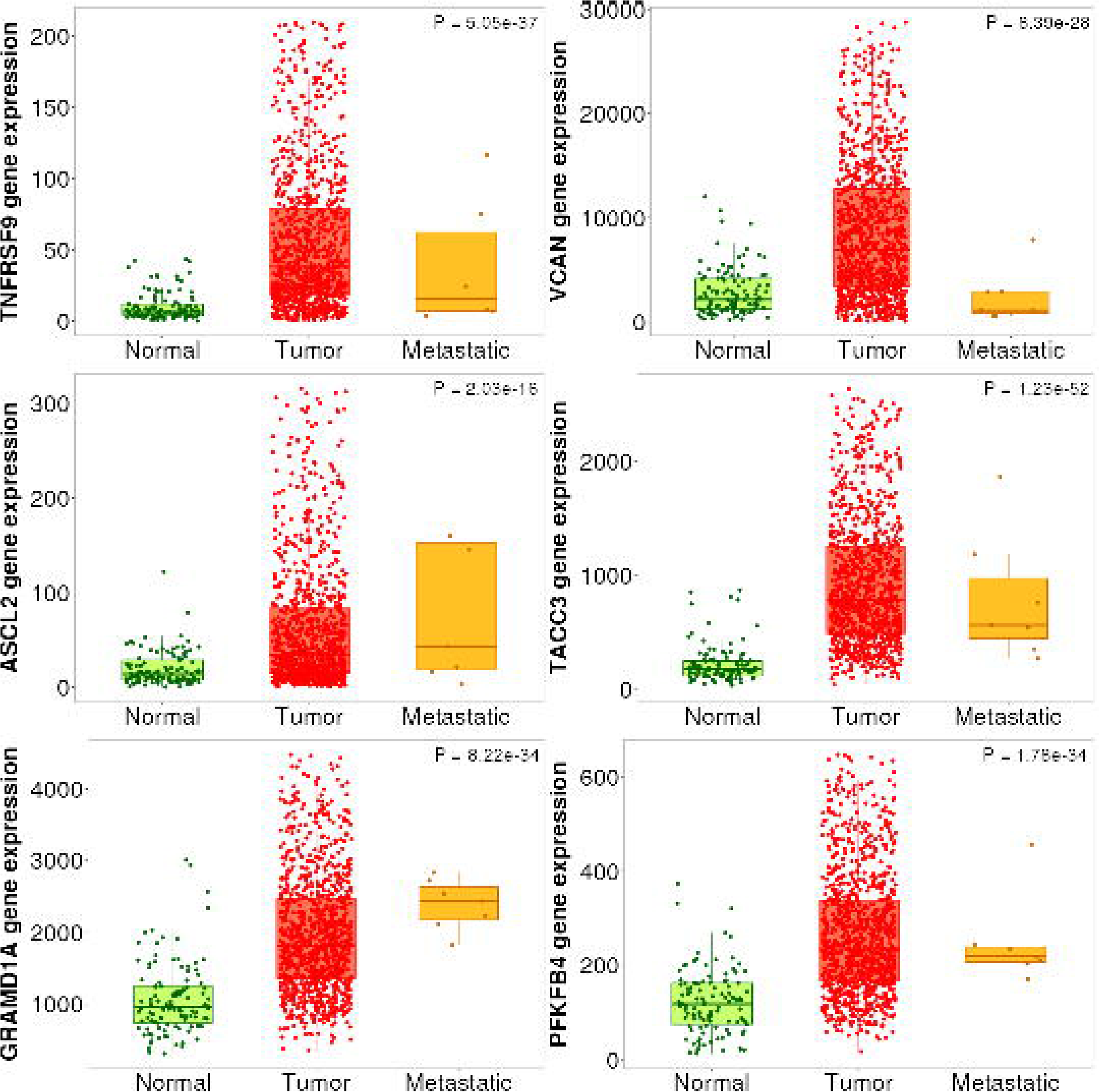
Gene expression profiles in breast cancer progression. Box plots show RNA-seq expression levels of *TNFRSF9*, *ASCL2*, *GRAMD1A*, *VCAN*, *TACC3*, and *PFKFB4* across normal breast tissue (n=113), primary tumors (n=1,097), and metastases (n=7). Kruskal-Wallis *P*-values (*P* < 1×10LJ¹LJ for all genes) confirm significant expression differences. *VCAN* is downregulated in metastases (*P* = 8.39×10LJ²LJ), while *TNFRSF9*, *ASCL2*, *TACC3*, *PFKFB4*, and *GRAMD1A* exhibit tumor-specific upregulation. Data source: TNMplot (GSE96058 cohort).

**Figure 4:**
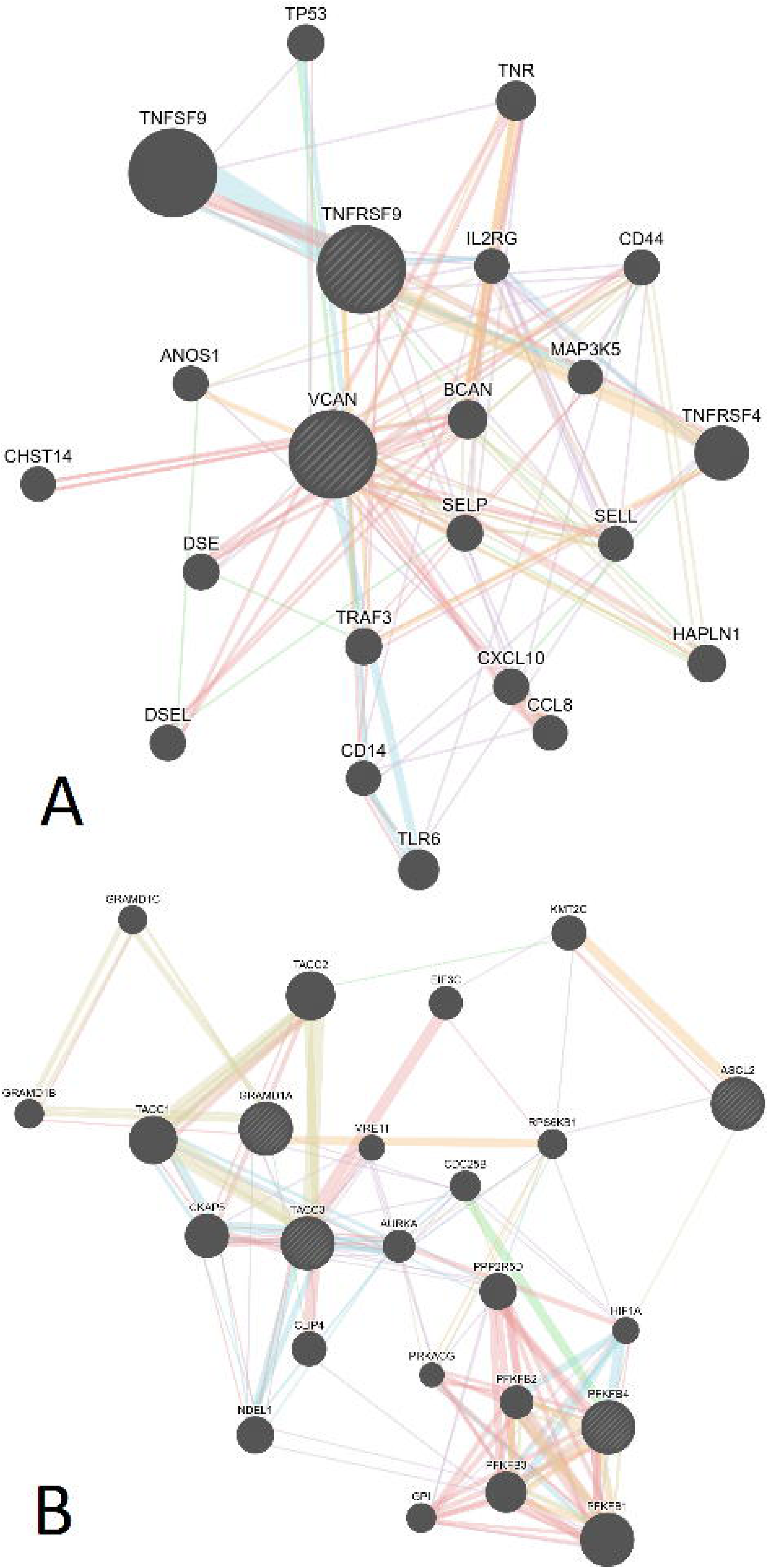
A | Downregulated gene interaction network. GeneMANIA-generated network showing functional associations for downregulated DEGs (*VCAN*, *CD44*, *TLR6*). Nodes represent genes; edges indicate co-expression (pink), physical interactions (blue), or shared pathways (yellow). Key clusters: immune signaling (*CXCL10*, *CCL8*), extracellular matrix remodeling (*VCAN*, *DSE*). **B | Upregulated gene interaction network.** GeneMANIA network for upregulated DEGs (*TACC3*, *PFKFB4*, *GRAMD1A*). Clusters include glycolysis (*PFKFB4*, *PFKFB1*), mitotic regulation (*TACC3*, *CKAP5*), and lipid metabolism (*GRAMD1A*). Edge colors denote interaction types as in panel A: co-expression (pink), physical interactions (blue), and shared pathways (yellow).

### Pathway Profiling

**Functional Enrichment**: Upregulated genes implicated NOTCH3-driven mitosis (TACC3), glycolysis (PFKFB4), and cholesterol metabolism (GRAMD1A). Downregulated genes tied to immune suppression (TNFRSF9) and stromal disengagement (VCAN) (Table 3).

**Table 3:** Functional Annotation of Upregulated and Downregulated DEGs in Brain Metastasis.

| GENE | KEY PATHWAYS | BIOLOGICAL PROCESSES (GO) | MOLECULAR FUNCTIONS (GO) | BIOLOGICAL RELEVANCE |
| --- | --- | --- | --- | --- |
| TACC3 | NOTCH3 activation (Reactome), RNA transport (KEGG), G2-M checkpoint (Hallmark) | Mitotic spindle organization, Cell cycle phase transition, Nuclear migration | --- | Links to mitotic fidelity and NOTCH3-driven stemness in metastasis |
| PFKFB4 | Glycolysis (Reactome), AMPK signaling (KEGG), Fructose metabolism (KEGG) | Organophosphate metabolic process, Response to nutrient levels | Phosphofructokinase activity, Sugar-phosphatase activity | Suggests metabolic reprogramming toward glycolysis in the brain microenvironment |
| ASCL2 | Neurogenesis (GO), Transcriptional regulation (GO) | Neuron differentiation, Placenta development, Negative regulation of transcription | RNA polymerase II repressor activity, E-box binding | Implicates neurodevelopmental plasticity in metastatic adaptation |
| GRAMD1A | Cholesterol transport (GO), Sterol metabolism (GO) | Cellular response to cholesterol, Intracellular lipid transport | Sterol binding, Cholesterol transfer activity | Indicates lipid metabolism rewiring for survival in cholesterol-rich brain tissue |
| VCAN | --- | Central nervous system development, Glial cell migration, Gliogenesis, Skeletal system development | Protein phosphatase activator activity | Downregulation suggests reduced involvement in nervous system development and glial cell migration, potentially impairing stromal interactions in the brain microenvironment |
| TNFRSF9 | TNFR2 non-canonical NF-κB pathway (Reactome), Cytokine-cytokine receptor interaction (KEGG) | Regulation of cell population proliferation | --- | Reduced cytokine signaling may indicate immune evasion or dampened pro-inflammatory responses in metastatic niches |

**Hub Gene-Centric Pathways**: Hub genes like VCAN enriched ECM-receptor interactions, while TNFRSF9 linked to T-cell signaling. PFKFB4 and TACC3 anchored hypoxia adaptation and mitotic control, respectively (Supplementary Figure 3 B&C).

### Common miRNAs Associated with Gene Networks

For the two downregulated DEGs (VCAN and TNFRSF9), miRNet predicted four miRNAs: hsa-miR-27a-3p, hsa-miR-210-3p, hsa-miR-1-3p, and hsa-miR-101-3p. Based on biological relevance to BCBM hallmarks (hypoxia, angiogenesis, metastasis), hsa-miR-27a-3p and hsa-miR-210-3p were prioritized. The latter two miRNAs (miR-1-3p and miR-101-3p) were excluded because their tumor-suppressive functions conflicted with the observed gene downregulation. For the upregulated oncogenes (PFKFB4, ASCL2, TACC3, GRAMD1A), miR-335 and miR-34a were identified as key regulators. (Supplementary Figure 4) Cross-referencing with **miRCancer** and PubMed validated these associations. (Supplementary Table 1)

In RNA-seq analysis of three datasets, MIR210HG (host gene for miR-210-3p) was significantly upregulated in GSE193103 (log2FC = 1.24, padj = 0.003) and showed a trend in GSE209998 (log2FC = 0.57, padj = 0.166), aligning with miR-210-3p’s role in hypoxia adaptation. But, MIR27A, MIR335, and MIR34A showed no significant differential expression (NS) or were absent in the datasetsPathway analysis linked these miRNAs to BCBM processes:

Pathway mapping via **miRPathDB** revealed that (Supplementary Table 2):

- **miR-27a-3p** and **miR-210-3p** drive hypoxia adaptation, metabolic reprogramming, and immune evasion, potentially suppressing *VCAN/TNFRSF9*.
- **miR-335** and **miR-34a** counteract oncogenic pathways (e.g., stemness, mitotic dysregulation) by targeting ECM remodeling and promoting apoptosis.

### Identification of Candidate Drugs Targeting Dysregulated Genes

Drug repurposing nominated Valproic Acid (epigenetic silencing of TACC3/ASCL2) and CDK4/6 inhibitors (Abemaciclib) for upregulated genes. For downregulated targets, HDAC inhibitors (Vorinostat) and chemotherapies (Cisplatin) emerged as candidates to restore VCAN/TNFRSF9 (Supplementary Table 3).

## Discussion

Our multi-omics framework identifies TNFRSF9 and VCAN as consistently suppressed in breast cancer brain metastasis (BCBM), alongside amplified expression of GRAMD1A, ASCL2, TACC3, and PFKFB4—driven by metastasis-specific genetic alterations. Elevated PFKFB4 and TACC3 in primary tumors predicted poor survival, while TNFRSF9/VCAN downregulation marked aggressive progression, positioning these genes as dual prognostic biomarkers and therapeutic targets. Pathway analyses uncovered BCBM mechanisms spanning extracellular matrix (ECM) remodeling, metabolic rewiring, and mitotic dysregulation. Upstream regulators (miR-210-3p, miR-34a) and repurposed drugs (Valproic Acid, CDK4/6 inhibitors) further highlighted actionable strategies. By bridging molecular drivers to clinical vulnerabilities, this integrative approach advances precision management of BCBM.

## TNFRSF9 in BCBM: Bridging Immune and Tumor-Suppressive Roles

Our findings on TNFRSF9 build upon its emerging role as a dual modulator of immune activation and tumor suppression. In melanoma, Fröhlich et al. (2020) linked TNFRSF9 hypomethylation to enhanced immune infiltration and anti-PD-1 response, [12] while Liu et al. (2022) demonstrated its tumor-suppressive function in breast cancer via p38/PAX6 signaling. [13] Our BCBM data bridge these themes: TNFRSF9 suppression in metastases— driven by deletions and hypoxia-responsive miRNAs (miR-27a-3p, miR-210-3p)—correlates with immune evasion and poor prognosis. Unlike prior studies focusing on intracellular signaling, we emphasize TNFRSF9’s immune-regulatory role in BCBM, mirroring its pan-cancer relevance. The identification of methotrexate and cisplatin as TNFRSF9-inducing agents aligns with therapeutic strategies to restore immune activation, offering a translational bridge for future studies.

## VCAN’s Paradox in Metastasis: Isoform-Specific Duality

The observed VCAN downregulation in BCBM contrasts with its well-documented pro-metastatic roles in ECM remodeling. [14–16] This paradox resolves through isoform-specific regulation: Sheng et al. showed the V1 isoform drives proliferation, while V2 suppresses metastasis. [17] Lee et al.’s work with amiodarone—shifting versican expression from V1 to V2—supports this dichotomy, inhibiting EMT in preclinical models. [18] Our findings align with Fanhchaksai et al. (2016), where stromal VCAN loss destabilizes ECM integrity, activating HA-CD44/TGFβ pathways to fuel metastasis. [19] Tissue-specific regulation is further underscored by Zhang et al. (2023), who tied VCAN suppression to aggressive breast phyllodes tumors. [20] These insights position VCAN as a context-dependent modulator, warranting isoform-resolved studies in BCBM.

## TACC3: A Recurrent Driver of Metastatic Adaptation

Our data reinforce TACC3’s role as a pan-cancer oncogene, altered (mainly amplified) in 23% of metastatic breast tumors versus 6% of primaries. Wang et al.’s meta-analysis (2017) [21] and Saatci and Sahin’s review (2023) [22] highlight TACC3’s ties to centrosome amplification, mitotic fidelity, and PI3K/AKT signaling—mechanisms echoed in our pathway analyses. Novel to this study is TACC3’s regulation by tumor-suppressive miRNAs (miR-34a, miR-335) and vulnerability to Valproic Acid or CDK4/6 inhibitors, offering clinically actionable strategies. These findings expand TACC3’s therapeutic relevance beyond FGFR3 fusions, positioning it as a central target in BCBM.

## PFKFB4: Metabolic Linchpin of Brain Metastasis

PFKFB4’s upregulation in BCBM (log2FC: +1.06–1.70) and poor prognostic impact (HR = 1.71) align with Dai et al.’s observations in TNBC metastases. [23] While prior work emphasized its role in redox balance via the pentose phosphate pathway, our data uniquely implicate PFKFB4 in AMPK/HIF-1 signaling, sustaining survival in the brain’s nutrient-poor niche. Network analysis further linked PFKFB4 to mTOR-driven cell cycle progression, diverging from Yao et al.’s focus on SRC3 interactions in luminal subtypes. [24] The prioritization of vorinostat and doxorubicin as glycolysis disruptors aligns with Kotowski et al.’s call [25] for metabolic targeting, underscoring PFKFB4’s therapeutic potential.

## ASCL2: Context-Dependent Architect of Metastatic Plasticity

ASCL2’s upregulation in BCBM (log2FC: +1.03–1.59) and ties to neurodevelopmental plasticity echo Xu et al. (2017) [26] and Faramarzy et al.’s work [27] on Wnt-driven stemness. However, its role diverges in neural malignancies—downregulated in brain tumors and melanoma—highlighting tissue-specific duality. Our finding of ASCL2 amplification (24% in metastases) and glucose metabolism roles expands its known Wnt/BIRC5 axis, suggesting niche-specific adaptations critical for brain colonization.

## GRAMD1A: Lipid Metabolism and Metastatic Resilience

GRAMD1A’s amplification (14% in metastases) and prognostic impact (HR = 1.45) position it as a mediator of lipid adaptation in BCBM. While Liu et al. [28] tied GRAMD1A to immune-metabolic crosstalk in renal cancer and Fu et al. to STAT5-driven stemness in HCC (29), our data emphasize lipid transport in the brain’s cholesterol-rich niche. Similarly, in Wilms tumor, GRAMD1A promotes proliferation and migration via cholesterol transport and autophagy (Zeng et al. [30]). Intriguingly, despite HCC links to doxorubicin resistance, we prioritized doxorubicin as a GRAMD1A suppressor, underscoring therapeutic contextuality.

## Strengths and Limitations

The study demonstrates notable strengths, including an integrative approach that harmonizes transcriptomic, genetic, and regulatory data across three cohorts, enhancing biological insight. Clinically, prognostic validation and drug prioritization (e.g., CDK4/6 inhibitors) directly link findings to therapeutic strategies, while pathway and network analyses reveal ECM remodeling, metabolic shifts, and mitotic drivers as key mechanisms. However, limitations exist: small metastatic sample sizes (e.g., seven in TNMplot) may reduce statistical reliability, and retrospective data introduce potential biases from batch effects or missing isoform-specific details (e.g., VCAN variants). Though prioritized miRNAs and drug candidates show promise, experimental validation remains critical to confirm their clinical relevance.

## Conclusions

This study systematically identifies TNFRSF9, VCAN, GRAMD1A, ASCL2, TACC3, and PFKFB4 as BCBM drivers, linking their dysregulation to genetic alterations, survival outcomes, and actionable pathways. By integrating multi-omics layers, we advance mechanistic insights into brain-specific adaptation and nominate clinically translatable therapies. These findings lay a foundation for precision strategies against BCBM, urging future studies to validate targets like CDK4/6 inhibitors and explore isoform-specific roles in metastasis.

## Acknowledgments

This article was reviewed and revised with the assistance of Depseek for linguistic editing, grammar correction, and text style optimization.

Researchers would like to express our sincere appreciation and our deepest gratitude to the Clinical Research Development Center of Kamali and Rajaee Hospitals in Alborz University of Medical Sciences.

