## Supplementary File for "Unveiling Novel Molecular Drivers in Breast Cancer Brain Metastasis: Multi-Omics Integration Identifies Downregulation of VCAN and Emerging Roles of ASCL2/GRAMD1A as Prognostic Biomarkers and Therapeutic Vulnerabilities"

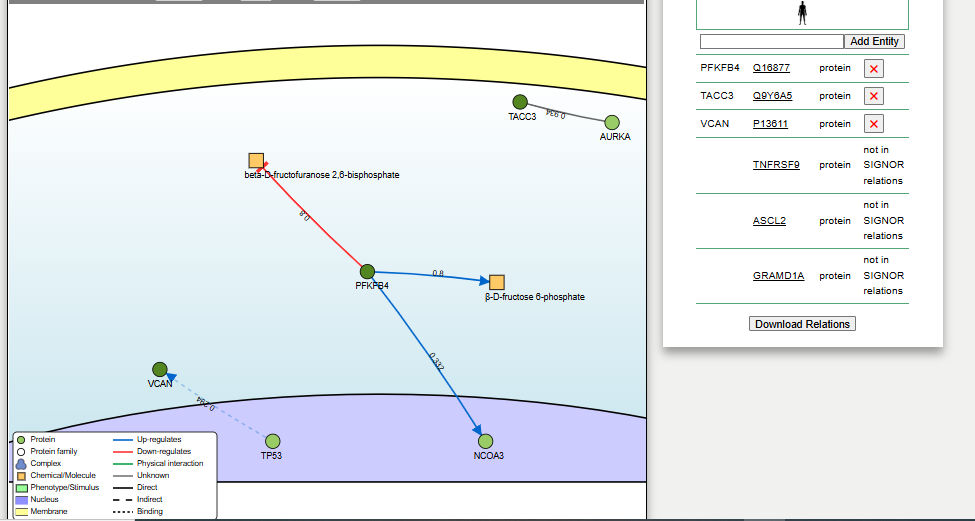

**Supplementary Figure 1. Signaling interactions of differentially expressed genes (DEGs) mapped to the SIGNOR database.**
Direct signaling interactions for DEGs were queried in the SIGNOR database ([https://signor.uniroma2.it](https://signor.uniroma2.it/)) using all available interaction scores. Arrows denote signaling relationships between genes. PFKFB4, TACC3, and VCAN were present in SIGNOR, while TNFRSF9, ASCL2, and GRAMD1A were absent, highlighting gaps in understanding their signaling roles

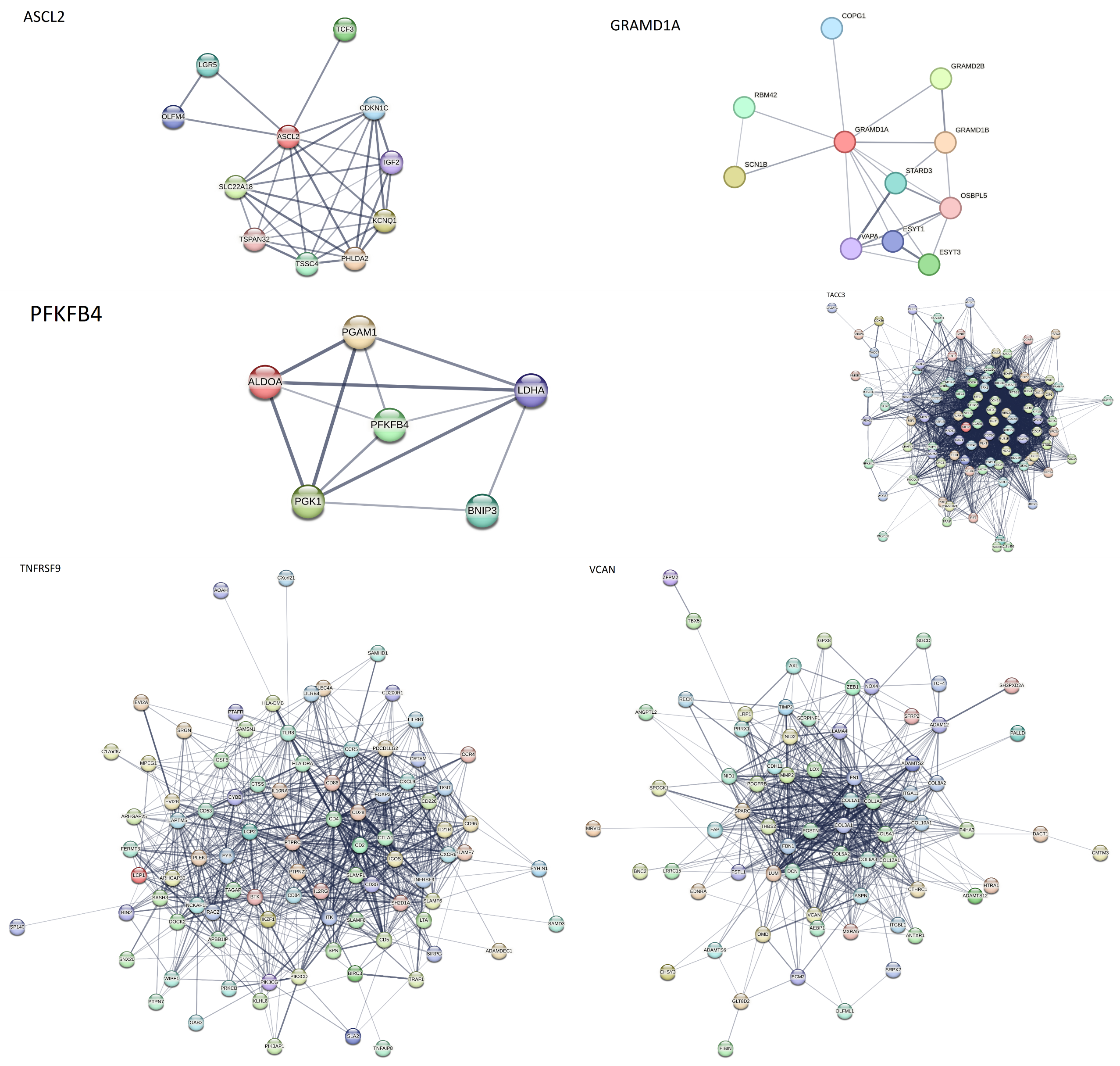

**Supplementary Figure 2.** A protein-protein interaction (PPI) network from the STRING database

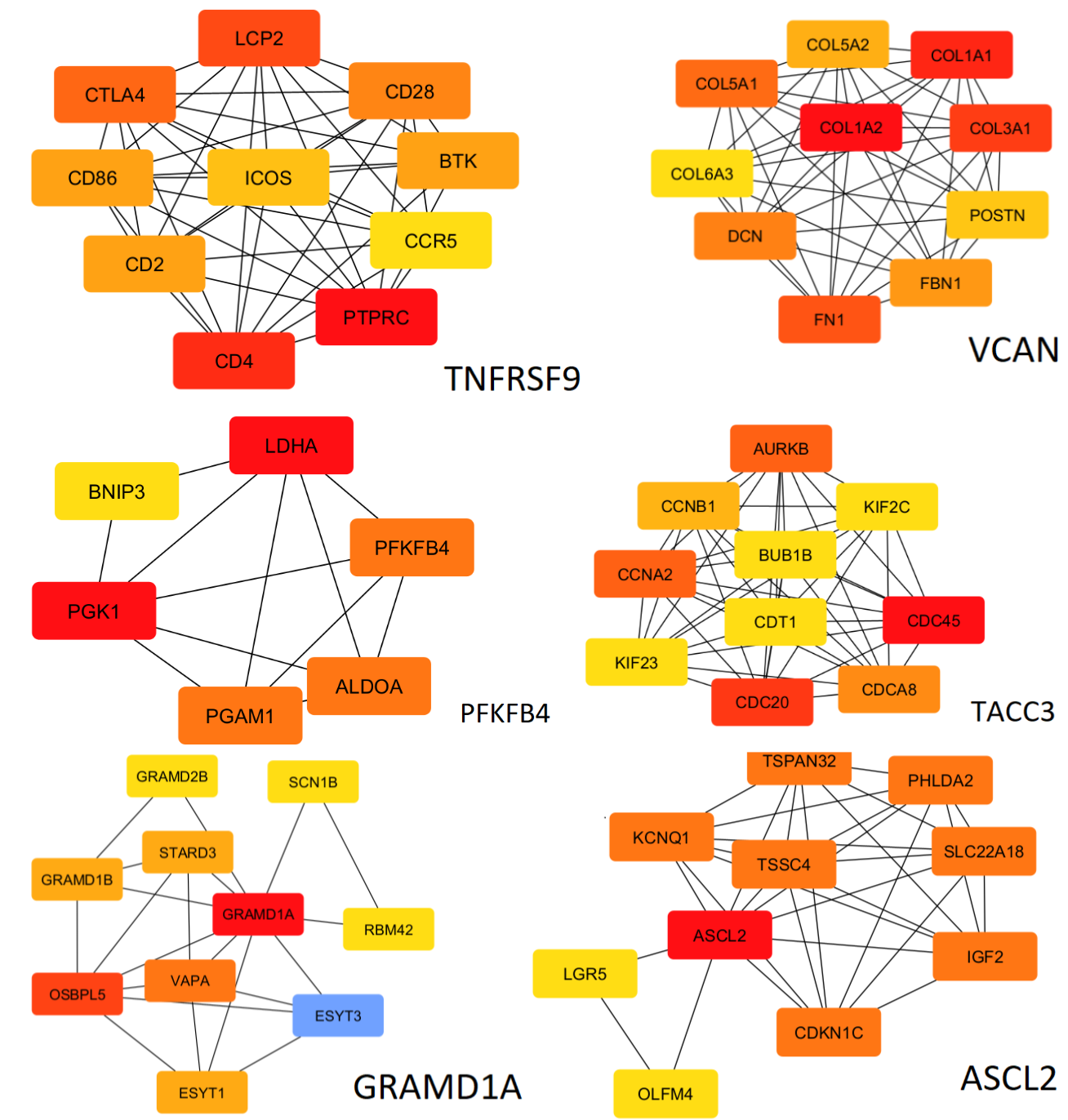

**A**

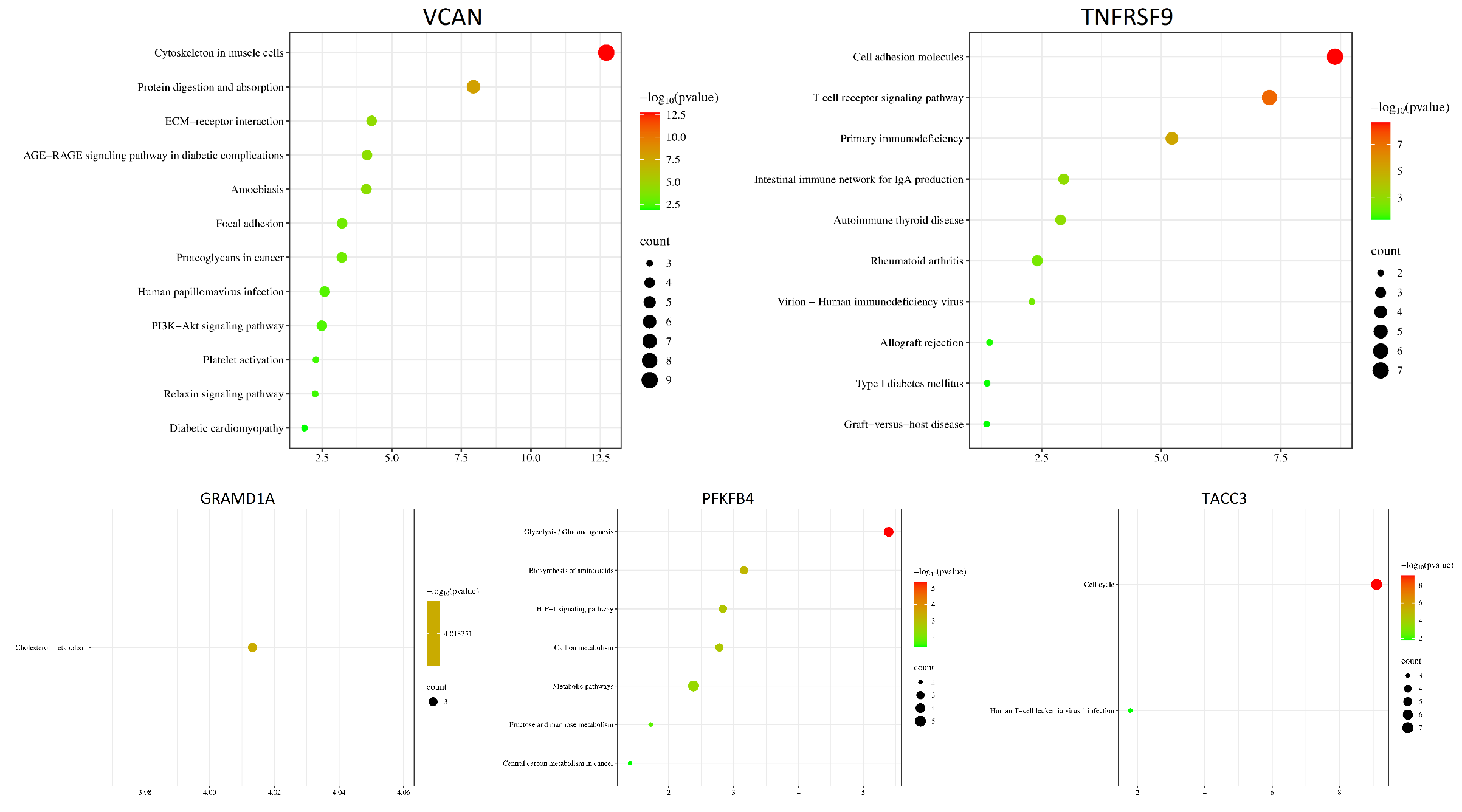

**B**

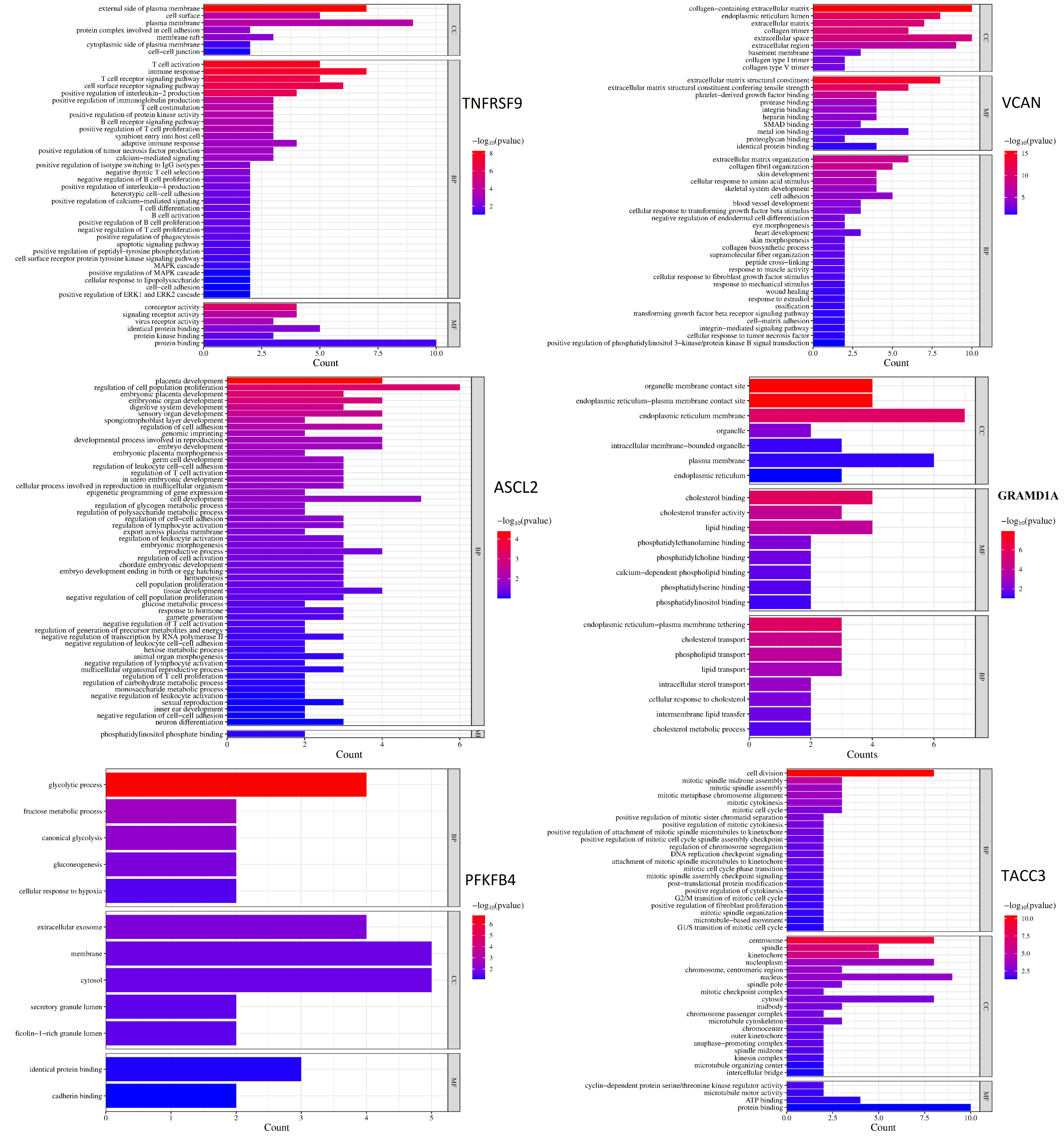

**C**

**Supplementary Figure 3.** Screening results of the functional analysis for key DEGs. **(A)** Top 10 hub genes prioritized by CytoHubba using the Degree algorithm, highlighting genes with highest network connectivity. **(B)** Enriched KEGG pathways for hub genes, including terms such as AGE-RAGE signaling (diabetic complications), T cell receptor signaling, and ECM-receptor interaction. **(C)** Gene Ontology (GO) enrichment across biological processes (BP), cellular components (CC), and molecular functions (MF). Pathways and terms were filtered by significance (−log _10_ (p-value)) and gene count.

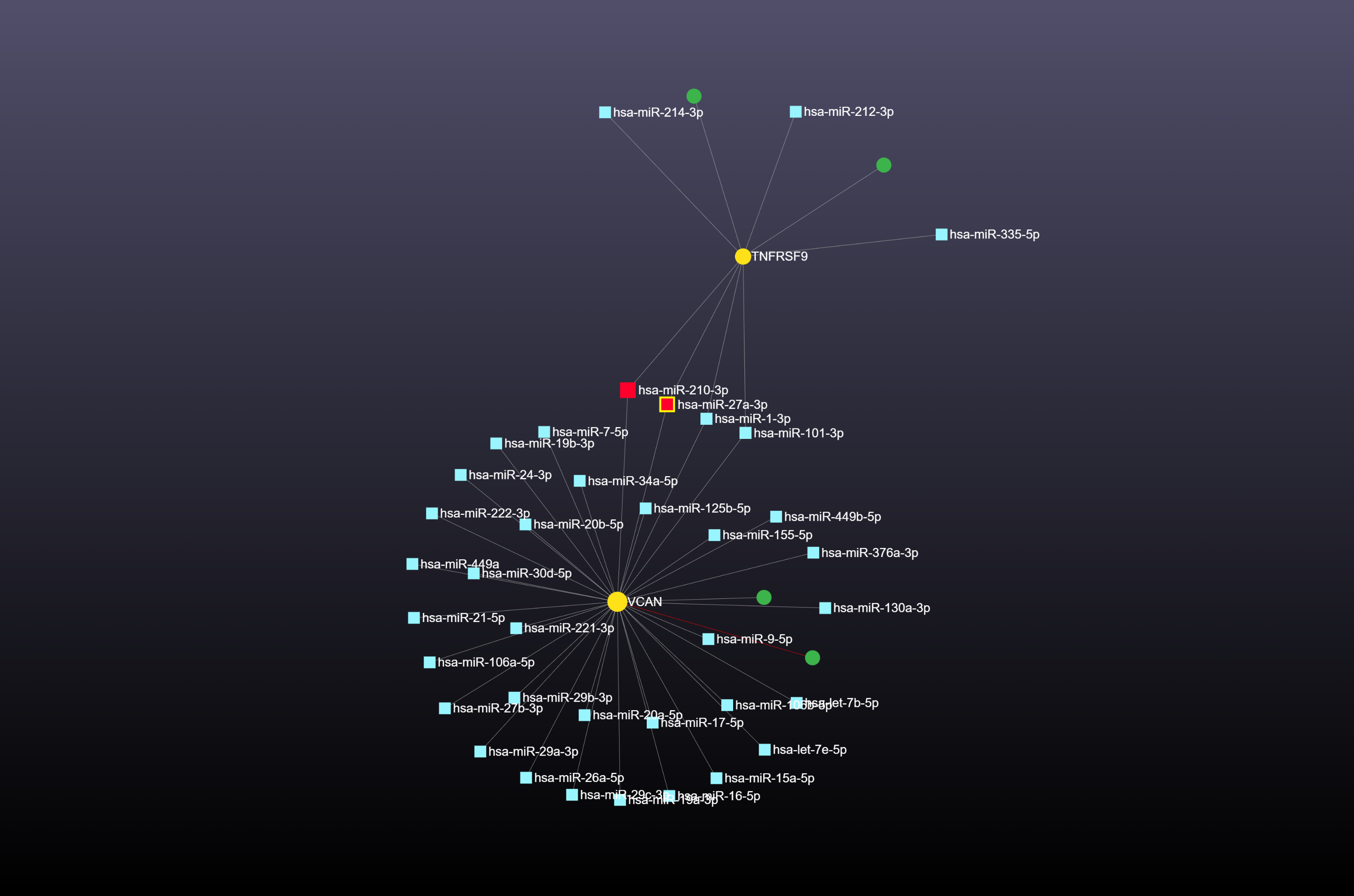
 **A**

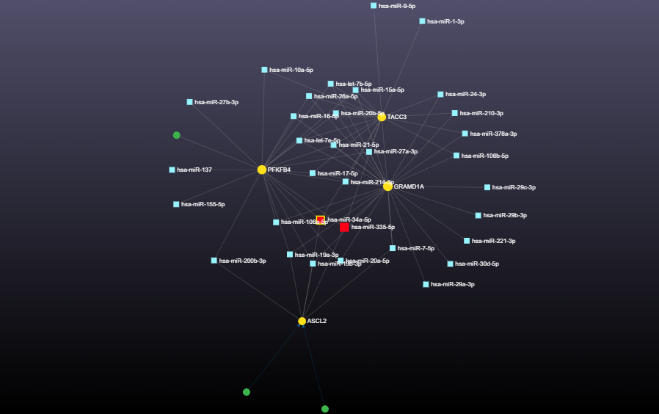
 **B**

**Supplementary Figure 4. Interaction network of miRNAs hsa-miR-27a-3p and hsa-miR-210-3p (yellow square nodes) targeting the down-regulated DEGs VCAN and TNFRSF9 (yellow circle nodes), generated using Mirnet. B: Network of miRNAs hsa-miR-335 and hsa-miR-34a (yellow square nodes) targeting the up-regulated oncogenes TACC3, GRAMD1A, PFKFB4, and ASCL2 (red circle nodes). The edges symbolize the interaction between the DEGs and related miRNAs**

**Supplementary Table 1: miRNAs Targeting DEGs:** Compiled PubMed-supported functional annotations (source: **mircancer.ecu.edu, March 2025)**

| Mir Id | Target | PubMed Article | More key findings | Confirming our findings |
| --- | --- | --- | --- | --- |
| hsa-miR-27a | VCAN TNFRSF9 | - c-MYC regulated miR-23a~24-2~27a cluster promotes mammary carcinoma cell invasion and hepatic metastasis by targeting Sprouty2. **(PMID: 23649631)** | **Upregulation of hsa-miR27a** drives breast cancer metastasis,  by suppressing the tumor suppressor SPRY2, activating MAPK signaling, and enhancing cell invasion and metastasis (as shown in hepatic metastasis) | + |
|  |  | - Arsenic trioxide suppresses cell growth and migration via inhibition of miR-27a in breast cancer cells. **(PMID: 26592661)** | miR-27a inhibitor treatment potentiates arsenic trioxide -induced breast cancer cell growth inhibition, apoptosis and motility inhibition | + |
|  |  | - miR-27a regulates the sensitivity of breast cancer cells to cisplatin treatment via BAK-SMAC/DIABLO-XIAP axis. **(PMID: 26662313)** | miR-27a is an onco-microRNA, promoteing the growth and metastasis of breast cancer cells.  the knockdown of miR-27a promoted the apoptosis via mitochondrial pathway in T-47D cells treated with  cisplatin (CDDP) | + |
| hsa-miR-210 | VCAN TNFRSF9 | - MicroRNA-210 interacts with FBXO31 to regulate cancer proliferation cell cycle and migration in human breast cancer. **(PMID: 27601917)** | miR-210 downregulation reduced cancer progression, induced cell cycle arrest, and inhibited cancer migration in T47D and MCF-7 cells. | + |
| hsa-miR-335 | PFKFB4, ASCL2, TACC3, GRAMD1A | MicroRNA-335 inhibits tumor reinitiation and is silenced through genetic and epigenetic mechanisms in human breast cancer. (PMID: 21289068) | miR-335 acts as a potent metastasis suppressor in breast cancer by inhibiting migration, invasion, and metastatic colonization. It targets genes like **SOX4** (a transcription factor) and **Tenascin-C (TNC)** (an extracellular matrix protein), which drive metastatic processes when miR-335 is downregulated. | + |
|  |  | Analysis of miR-205 and miR-155 expression in the blood of breast cancer patients. (PMID: 23372341) | - Restoring miR-335 could suppress metastasis and tumor growth.  -low miR-335 levels correlate with shorter median time to metastatic relapse in BC patients(Ref.: Tavazoie et al., 2008) | + |
|  |  | - MiR-335 inhibits migration of breast cancer cells through targeting oncoprotein c-Met. (PMID: 25492484) | miR-335 suppresses breast cancer cell migration by negatively regulating the HGF/c-Met pathway | + |
|  |  | - MicroRNA-335 suppresses the proliferation, migration, and invasion of breast cancer cells by targeting EphA4. (PMID: 28795314) | - | + |
| hsa-miR-34a | PFKFB4, ASCL2, TACC3, GRAMD1A | - MicroRNA-34 suppresses breast cancer invasion and metastasis by directly targeting Fra-1. (PMID: 23001043) | - | + |
|  |  | - Targeted Expression of miR-34a Using the T-VISA System Suppresses Breast Cancer Cell Growth and Invasion. (PMID: 23032974) | - | + |
|  |  | - Tumor-suppressive microRNA-34a inhibits breast cancer cell migration and invasion via targeting oncogenic TPD52. (PMID: 26678891) | - | + |
|  |  | - MiR-34a Inhibits Breast Cancer Proliferation and Progression by Targeting Wnt1 in Wnt/?-Catenin Signaling Pathway. (PMID: 27524218) | - | + |
|  |  | miR-34a expression in human breast cancer is associated with drug resistance. ( PMID: 29290947) | - | + |
|  |  | MicroRNA-34a suppresses breast cancer cell proliferation and invasion by targeting Notch1. (PMID: 30542388) | - | + |

**Supplementary Table 2:** Pathway mapping via **miRPathDB**

| miRNA | Database | Pathway/Process | Evidence | P-value | Rationale |
| --- | --- | --- | --- | --- | --- |
| miR-34a | KEGG | Colorectal cancer | experimental (strong) | 1.96E-06 | Direct links to miR-34a’s role in tumor suppression via p53 activation, Wnt inhibition, and oncogene silencing. |
| miR-34a | KEGG | MicroRNAs in cancer | experimental (strong) | 1.32E-05 | Direct links to miR-34a’s role in tumor suppression via p53 activation, Wnt inhibition, and oncogene silencing. |
| miR-34a | KEGG | p53 signaling pathway | experimental (strong) | 0.017 | Direct links to miR-34a’s role in tumor suppression via p53 activation, Wnt inhibition, and oncogene silencing. |
| miR-34a | KEGG | Wnt signaling pathway | experimental (strong) | 0.019 | Direct links to miR-34a’s role in tumor suppression via p53 activation, Wnt inhibition, and oncogene silencing. |
| miR-34a | WikiPathways | Breast cancer pathway | experimental (strong) | 2.16E-06 | EMT inhibition (via SNAIL repression), TGF-β signaling, and hypoxia adaptation block metastasis. |
| miR-34a | WikiPathways | Epithelial to mesenchymal transition in colorectal cancer | experimental (strong) | 0.007 | EMT inhibition (via SNAIL repression), TGF-β signaling, and hypoxia adaptation block metastasis. |
| miR-34a | WikiPathways | TGF-beta Signaling Pathway | experimental (strong) | 0.033 | EMT inhibition (via SNAIL repression), TGF-β signaling, and hypoxia adaptation block metastasis. |
| miR-34a | WikiPathways | HIF-1 signaling pathway | experimental (strong) | 0.047 | EMT inhibition (via SNAIL repression), TGF-β signaling, and hypoxia adaptation block metastasis. |
| miR-34a | Reactome | Signaling by NOTCH | experimental (strong) | 0.002 | NOTCH and PI3K-AKT are pro-metastatic pathways targeted by miR-34a. Autophagy supports survival under stress. |
| miR-34a | Reactome | PI3K/AKT signaling | experimental (strong) | 0.035 | NOTCH and PI3K-AKT are pro-metastatic pathways targeted by miR-34a. Autophagy supports survival under stress. |
| miR-34a | Reactome | Autophagy | experimental (strong) | 0.005 | NOTCH and PI3K-AKT are pro-metastatic pathways targeted by miR-34a. Autophagy supports survival under stress. |
| miR-34a | Reactome | PIP3 activates AKT signaling | experimental (strong) | 0.035 | NOTCH and PI3K-AKT are pro-metastatic pathways targeted by miR-34a. Autophagy supports survival under stress. |
| miR-34a | Gene Ontology - Biological Process | apoptotic process | experimental (strong) | 0.006 | miR-34a induces apoptosis, inhibits proliferation, and suppresses Wnt/β-catenin signaling to block metastasis. |
| miR-34a | Gene Ontology - Biological Process | negative regulation of cell death | experimental (strong) | 0.003 | miR-34a induces apoptosis, inhibits proliferation, and suppresses Wnt/β-catenin signaling to block metastasis. |
| miR-34a | Gene Ontology - Biological Process | regulation of cell population proliferation | experimental (strong) | 0.002 | miR-34a induces apoptosis, inhibits proliferation, and suppresses Wnt/β-catenin signaling to block metastasis. |
| miR-34a | Gene Ontology - Biological Process | Wnt signaling pathway | experimental (strong) | 0.019 | miR-34a induces apoptosis, inhibits proliferation, and suppresses Wnt/β-catenin signaling to block metastasis. |
| miR-34a | Gene Ontology - Biological Process | response to hypoxia | experimental (strong) | 0.014 | Hypoxia and oxidative stress resistance are hallmarks of metastatic cells; miR-34a modulates these pathways. |
| miR-34a | Gene Ontology - Biological Process | cellular response to oxidative stress | experimental (strong) | 0.004 | Hypoxia and oxidative stress resistance are hallmarks of metastatic cells; miR-34a modulates these pathways. |
| miR-34a | Gene Ontology - Biological Process | regulation of autophagy | experimental (strong) | 0.003 | Hypoxia and oxidative stress resistance are hallmarks of metastatic cells; miR-34a modulates these pathways. |
| miR-335 | Gene Ontology - Cellular Component | extracellular matrix | experimental (any) | 3.01E-06 | miR-335 targets ECM components (e.g., collagen), disrupting matrix integrity and hindering tumor invasion. |
| miR-335 | Gene Ontology - Cellular Component | collagen-containing extracellular matrix | experimental (any) | 1.89E-05 | miR-335 targets ECM components (e.g., collagen), disrupting matrix integrity and hindering tumor invasion. |
| miR-335 | Gene Ontology - Molecular Function | extracellular matrix structural constituent | experimental (any) | 8.71E-04 | Binds ECM structural proteins to limit matrix stiffness and blocks receptor signaling critical for migration. |
| miR-335 | Gene Ontology - Molecular Function | signaling receptor activity | experimental (any) | 4.65E-12 | Binds ECM structural proteins to limit matrix stiffness and blocks receptor signaling critical for migration. |
| miR-335 | Reactome | Extracellular matrix organization | experimental (any) | 0.008 | Suppresses ECM remodeling (e.g., collagen degradation) and inhibits GPCR/chemokine-driven migration. |
| miR-335 | Reactome | Signaling by GPCR | experimental (any) | 9.64E-05 | Suppresses ECM remodeling (e.g., collagen degradation) and inhibits GPCR/chemokine-driven migration. |
| miR-335 | Reactome | Chemokine receptors bind chemokines | experimental (any) | 0.006 | Suppresses ECM remodeling (e.g., collagen degradation) and inhibits GPCR/chemokine-driven migration. |
| miR-335 | KEGG | Cytokine-cytokine receptor interaction | experimental (any) | 0.003 | Modulates cytokine signaling (e.g., IL-6/STAT3) to reduce pro-metastatic inflammation. |
| miR-335 | KEGG | Complement and coagulation cascades | experimental (any) | 0.012 | Modulates cytokine signaling (e.g., IL-6/STAT3) to reduce pro-metastatic inflammation. |
| miR-335 | KEGG | MicroRNAs in cancer | experimental (strong) | 0.003 | Modulates cytokine signaling (e.g., IL-6/STAT3) to reduce pro-metastatic inflammation. |
| miR-335 | WikiPathways | TGF-beta Signaling Pathway | experimental (strong) | 0.01 | Inhibits TGF-β-induced EMT and metastasis in breast cancer by targeting SMAD2/3 or other effectors. |
| miR-335 | WikiPathways | Breast cancer pathway | experimental (strong) | 0.017 | Inhibits TGF-β-induced EMT and metastasis in breast cancer by targeting SMAD2/3 or other effectors. |
| miR-335 | Gene Ontology - Biological Process | cell migration | experimental (any) | 0.012 | Directly suppresses migration (via RHOA/ROCK inhibition) and angiogenesis. |
| miR-335 | Gene Ontology - Biological Process | angiogenesis | experimental (any) | 0.01 | Directly suppresses migration (via RHOA/ROCK inhibition) and angiogenesis. |
| miR-335 | Gene Ontology - Biological Process | regulation of cell motility | experimental (any) | 0.039 | Directly suppresses migration (via RHOA/ROCK inhibition) and angiogenesis. |
| miR-210 | Gene Ontology - Biological Process | cellular response to hypoxia | experimental (any) | 0.01 | Hypoxia drives metastasis by promoting angiogenesis, invasion, and EMT. |
| miR-210 | Gene Ontology - Biological Process | cellular response to oxygen levels | experimental (any) | 0.007 | Hypoxia drives metastasis by promoting angiogenesis, invasion, and EMT. |
| miR-210 | Gene Ontology - Biological Process | response to oxidative stress | experimental (any) | 0.025 | Hypoxia drives metastasis by promoting angiogenesis, invasion, and EMT. |
| miR-210 | Gene Ontology - Biological Process | regulation of cell differentiation | experimental (any) | 0.007 | EMT and dedifferentiation are critical for metastatic spread. |
| miR-210 | Gene Ontology - Biological Process | negative regulation of gene expression | experimental (any) | 0.008 | EMT and dedifferentiation are critical for metastatic spread. |
| miR-210 | Gene Ontology - Biological Process | cell migration | experimental (any) | 0.012 | EMT and dedifferentiation are critical for metastatic spread. |
| miR-210 | KEGG | Transcriptional misregulation in cancer | experimental (any) | 0.015 | Direct link to oncogenic transcriptional dysregulation, including miRNA roles in metastasis. |
| miR-210 | WikiPathways | Cell migration and invasion through p75NTR | experimental (any) | 0.031 | p75NTR promotes invasion; HIF-1α stabilizes under hypoxia to drive glycolysis and metastasis. |
| miR-210 | WikiPathways | HIF1A and PPARG regulation of glycolysis | experimental (any) | 0.031 | p75NTR promotes invasion; HIF-1α stabilizes under hypoxia to drive glycolysis and metastasis. |
| miR-210 | Gene Ontology - Biological Process | apoptotic process | experimental (any) | 0.016 | Evasion of apoptosis is a hallmark of metastatic cells surviving oxidative stress. |
| miR-210 | Gene Ontology - Biological Process | intrinsic apoptotic signaling pathway in response to oxidative stress | experimental (any) | 0.044 | Evasion of apoptosis is a hallmark of metastatic cells surviving oxidative stress. |
| miR-210 | Gene Ontology - Biological Process | oxidation-reduction process | experimental (strong) | 0.043 | Metabolic reprogramming (Warburg effect) fuels metastasis. Strong experimental evidence. |
| miR-210 | Gene Ontology - Biological Process | aerobic respiration | experimental (strong) | 0.043 | Metabolic reprogramming (Warburg effect) fuels metastasis. Strong experimental evidence. |
| miR-210 | Reactome | Pyruvate metabolism and Citric Acid (TCA) cycle | experimental (strong) | 0.048 | Mitochondrial metabolism supports metastatic cell survival. |
| miR-210 | Reactome | Respiratory electron transport | experimental (strong) | 0.048 | Mitochondrial metabolism supports metastatic cell survival. |
| miR-210 | Gene Ontology - Molecular Function | oxidoreductase activity | experimental (strong) | 0.031 | Lactate production (via LDH) acidifies microenvironment, promoting invasion. |
| miR-210 | Gene Ontology - Molecular Function | L-lactate dehydrogenase activity | experimental (strong) | 0.031 | Lactate production (via LDH) acidifies microenvironment, promoting invasion. |
| miR-210 | Gene Ontology - Biological Process | regulation of angiogenesis | experimental (any) | 0.034 | Angiogenesis is essential for nutrient supply to metastatic tumors. |
| miR-210 | Gene Ontology - Biological Process | vasculature development | experimental (any) | 0.043 | Angiogenesis is essential for nutrient supply to metastatic tumors. |
| miR-210 | Gene Ontology - Biological Process | negative regulation of microtubule depolymerization | experimental (any) | 0.008 | Microtubule dynamics regulate cell motility and invasion. |
| miR-27a | KEGG | MicroRNAs in cancer | experimental (strong) | 0.041 | Direct links to miRNA-driven oncogenesis. PI3K-AKT promotes survival/metastasis. |
| miR-27a | KEGG | PI3K-Akt signaling pathway | experimental (strong) | 0.002 | Direct links to miRNA-driven oncogenesis. PI3K-AKT promotes survival/metastasis. |
| miR-27a | KEGG | Transcriptional misregulation in cancer | experimental (strong) | 0.006 | Direct links to miRNA-driven oncogenesis. PI3K-AKT promotes survival/metastasis. |
| miR-27a | WikiPathways | Integrated Breast Cancer Pathway | experimental (strong) | 8.76E-06 | EMT is critical for invasion. TGF-β drives stromal interactions and metastasis. |
| miR-27a | WikiPathways | Epithelial to mesenchymal transition in colorectal cancer | experimental (strong) | 0.007 | EMT is critical for invasion. TGF-β drives stromal interactions and metastasis. |
| miR-27a | WikiPathways | TGF-beta Signaling Pathway | experimental (strong) | 0.002 | EMT is critical for invasion. TGF-β drives stromal interactions and metastasis. |
| miR-27a | Reactome | Signaling by EGFR | experimental (strong) | 0.047 | EGFR/PI3K-AKT are central to migration and survival. SUMOylation modifies HIF-1α for hypoxic adaptation. |
| miR-27a | Reactome | PI3K-AKT signaling | experimental (strong) | 0.047 | EGFR/PI3K-AKT are central to migration and survival. SUMOylation modifies HIF-1α for hypoxic adaptation. |
| miR-27a | Reactome | SUMOylation | experimental (strong) | 0.029 | EGFR/PI3K-AKT are central to migration and survival. SUMOylation modifies HIF-1α for hypoxic adaptation. |
| miR-27a | Gene Ontology - Biological Process | cell migration | experimental (strong) | 0.012 | Direct roles in metastasis: proliferation sustains tumor growth; angiogenesis fuels nutrient supply. |
| miR-27a | Gene Ontology - Biological Process | positive regulation of cell population proliferation | experimental (strong) | 0.046 | Direct roles in metastasis: proliferation sustains tumor growth; angiogenesis fuels nutrient supply. |
| miR-27a | Gene Ontology - Biological Process | regulation of angiogenesis | experimental (strong) | 0.023 | Direct roles in metastasis: proliferation sustains tumor growth; angiogenesis fuels nutrient supply. |
| miR-27a | Gene Ontology - Biological Process | apoptotic process | experimental (strong) | 0.006 | Evasion of apoptosis and hypoxia adaptation are hallmarks of metastatic cells. |
| miR-27a | Gene Ontology - Biological Process | negative regulation of cell death | experimental (strong) | 7.83E-04 | Evasion of apoptosis and hypoxia adaptation are hallmarks of metastatic cells. |
| miR-27a | Gene Ontology - Biological Process | response to hypoxia | experimental (strong) | 0.021 | Evasion of apoptosis and hypoxia adaptation are hallmarks of metastatic cells. |
| miR-27a | Gene Ontology - Molecular Function | phosphatase binding | experimental (strong) | 2.01E-04 | miR-27a may target phosphatases (e.g., PTEN) to activate PI3K-AKT. |
| miR-27a | Gene Ontology - Molecular Function | DNA-binding transcription factor activity | experimental (strong) | 0.005 | miR-27a may target phosphatases (e.g., PTEN) to activate PI3K-AKT. |
| miR-27a | Gene Ontology - Molecular Function | protein kinase activity | experimental (strong) | 0.015 | miR-27a may target phosphatases (e.g., PTEN) to activate PI3K-AKT. |
| miR-27a | WikiPathways | Cell migration and invasion through p75NTR | experimental (strong) | 0.031 | p75NTR promotes invasion; HIF-1α drives glycolysis under hypoxia. |
| miR-27a | WikiPathways | HIF1A and PPARG regulation of glycolysis | experimental (strong) | 0.031 | p75NTR promotes invasion; HIF-1α drives glycolysis under hypoxia. |
| miR-27a | KEGG | Prostate cancer | experimental (strong) | 6.58E-04 | Tissue-specific pathways where miR-27a is implicated in metastasis. |
| miR-27a | KEGG | Colorectal cancer | experimental (strong) | 5.27E-04 | Tissue-specific pathways where miR-27a is implicated in metastasis. |
| miR-27a | KEGG | Thyroid cancer | experimental (strong) | 0.001 | Tissue-specific pathways where miR-27a is implicated in metastasis. |
| miR-27a | Gene Ontology - Biological Process | Wnt signaling pathway | experimental (strong) | 0.019 | Wnt signaling drives EMT and stemness. Differentiation regulation impacts metastatic plasticity. |
| miR-27a | Gene Ontology - Biological Process | regulation of cell differentiation | experimental (strong) | 0.019 | Wnt signaling drives EMT and stemness. Differentiation regulation impacts metastatic plasticity. |
| miR-27a | Reactome | Hemostasis | experimental (strong) | 0.011 | Platelets shield circulating tumor cells, aiding immune evasion and extravasation. |
| miR-27a | Reactome | Platelet activation | experimental (strong) | 0.047 | Platelets shield circulating tumor cells, aiding immune evasion and extravasation. |

**Supplementary Table 3:** Candidate Drugs Targeting DEGs

| GENE | REGULATION | CANDIDATE DRUGS | MECHANISM | CLINICAL RELEVANCE |
| --- | --- | --- | --- | --- |
| TACC3 | Upregulated | Valproic Acid, Abemaciclib | Epigenetic silencing; CDK4/6 inhibition | FDA-approved for breast cancer |
| ASCL2 | Upregulated | Valproic Acid, Tretinoin | Methylation; differentiation induction | Epigenetic/APL therapy |
| PFKFB4 | Upregulated | Vorinostat, Doxorubicin | HDAC inhibition; glycolysis blockade | Lymphoma/breast cancer chemo |
| TNFRSF9 | Downregulated | Methotrexate, Cisplatin | Immune activation via mRNA upregulation | Standard solid tumor chemo |
| VCAN | Downregulated | Valproic Acid, Vorinostat | Epigenetic restoration of expression | BBB-penetrant HDAC inhibitors |
